# Epigallocatechin Gallate from Green Tea Effectively Blocks Infection of SARS-CoV-2 and New Variants by Inhibiting Spike Binding to ACE2 Receptor

**DOI:** 10.1101/2021.03.17.435637

**Authors:** Jinbiao Liu, Brittany H Bodnar, Fengzhen Meng, Adil Khan, Xu Wang, Guangxiang Luo, Sami Saribas, Tao Wang, Saroj Chandra Lohani, Peng Wang, Zhengyu Wei, Jinjun Luo, Lina Zhou, Jianguo Wu, Qingsheng Li, Wenhui Hu, Wenzhe Ho

## Abstract

As the COVID-19 pandemic rages on, the new SARS-CoV-2 variants have emerged in the different regions of the world. These newly emerged variants have mutations in their spike (S) protein that may confer resistance to vaccine-elicited immunity and existing neutralizing antibody therapeutics. Therefore, there is still an urgent need of safe, effective, and affordable agents for prevention/treatment of SARS-CoV-2 and its variant infection. Here, we demonstrated that green tea beverage (GTB) or its major ingredient, epigallocatechin gallate (EGCG), were highly effective in inhibiting infection of live SARS-CoV-2 and human coronavirus (HCoV OC43). In addition, infection of the pseudoviruses with spikes of the new variants (UK-B.1.1.7, SA-B.1.351, and CA-B.1.429) was efficiently blocked by GTB or EGCG. Among the 4 active green tea catechins at noncytotoxic doses, EGCG was the most potent in the action against the viruses. The highest inhibitory activity was observed when the viruses or the cells were pre-incubated with EGCG prior to the infection. Mechanistic studies revealed that EGCG blocked infection at the entry step through interfering with the engagement of the receptor binding domain (RBD) of the viral spikes to angiotensin-converting enzyme 2 (ACE2) receptor of the host cells. These data support further clinical evaluation and development of EGCG as a novel, safe, and cost-effective natural product for prevention/treatment of SARS-CoV-2 transmission and infection.

## Introduction

Severe acute respiratory syndrome coronavirus-2 (SARS-CoV-2), the causative agent of the pandemic coronavirus disease 2019 (COVID-19), has resulted in millions of deaths and more morbidities worldwide. Worst of all, as the COVID-19 pandemic rages on, there have emerged variants of SARS-CoV-2 in the different regions of the world, including the United Kingdom (UK), South Africa (SA), Brazil, and the United States^[1-3]^. These new variants have multiple mutations in their spike (S) protein, the key glycoprotein for SARS-CoV-2 infection, host immune response and antiviral therapy. Emerging of the SARS-CoV-2 with mutant S has sparked a great concern of whether the current COVID-19 vaccines are effective against the new variants. It has been demonstrated that these variants with increased infectivity may escape vaccine-induced immunity against COVID-19 and resist existing therapeutics^[4]^. Therefore, there is still an urgent need of safe, effective, and affordable prevention/treatment agents for SARS-CoV-2 infection.

SARS-CoV-2 infection is initiated when the receptor binding domain (RBD) of S protein binds to angiotensin-converting enzyme 2 (ACE2) receptor on the surface of host cells^[5]^. Because of the key role of S protein in SARS-CoV-2 infection, intervention with highly potent and cost-effective inhibitors that target at entry step of SARS-CoV-2 infection is an ideal strategy for slowing the COVID-19 pandemic. To date, there are no highly effective therapeutics available for blocking SARS-CoV-2 infection, except the neutralizing monoclonal antibodies, despite various approaches targeting SARS-CoV-2 entry have been attempted including the inhibitors of the viral S, as well as soluble ACE2^[6-10]^.

Green tea is one of the most widely consumed beverages in the world, particularly in Asian countries where the morbidity and mortality of COVID-19 are relatively low. Green tea is known to have many physiological and pharmacological health benefits^[11]^. The major active ingredients in green tea are the catechins, including (−)-epigallocatechin gallate (EGCG), (−)-epigallocatechin (EGC), (−)-epicatechin gallate (ECG) and (−)-epicatechin (EC). EGCG is the most abundant catechin in green tea^[12]^. Because of its unique polyphenolic structure, EGCG possesses antioxidant and anti-inflammatory properties^[12, 13]^. In addition, EGCG has been reported to have a broad-spectrum antiviral effect on several pathologic human viruses^[14-20]^, including those causing respiratory diseases^[21]^. Previous *in vitro* and *in vivo* studies ^[22-24]^ have shown that the green tea catechins could inhibit adsorption, replication, and neuraminidase activity of influenza A virus. A recent study^[25]^ demonstrated that the green tea catechins adsorbed on the murine pharyngeal mucosa reduced influenza A virus infection. In addition, the epidemiological studies from randomized controlled trials suggested that the regular consumption of green tea decreased influenza infection rates^[26, 27]^. Furthermore, results from a placebo-controlled and single-blind study showed that consumption of green tea catechins-containing beverage could prevent acute upper respiratory tract infections^[28]^.

Accumulating evidence indicates that green tea and its major ingredients (catechins) are beneficial for fighting COVID-19^[29]^. Green tea catechins are known to have impact on the factors associated with COVID-19 mortality, including antioxidative, anti-hypertensive, anti-proliferative, anti-thrombogenic, and lipid/cholesterol-lowering effects^[30]^. EGCG’s anti-inflammatory property may reduce overactivity of the immune system and ameliorate severity of COVID-19^[31]^. A recent *in vitro* study showed that EGCG can inhibit SARS-CoV-2 3CL-protease, an important enzyme for coronavirus infection/replication in the host^[32]^. These observations indicate that green tea could directly and indirectly reduce overall risks related to COVID-19. This assumption is further supported by a most recent ecological study^[30]^ showing that countries with higher level of green tea consumption had less morbidity and mortality of COVID-19, suggesting a possible etiological correlation between green tea consumption and incidence of SARS-CoV-2 infection. Therefore, there is a need of more experimental and mechanistic investigations to directly evaluate the impact of green tea and its major ingredients on SARS-CoV-2 infection. In this study, we examined whether green tea beverage (GTB) and the tea catechins (EGCG, EGC, ECG and EC) can inhibit infection of live SARS-CoV-2 and the pseudoviruses with spikes of the newly emerged variants (UK-B.1.1.7, SA-B.1.351, and CA-B.1.429). We also determined whether EGCG can block SARS-CoV-2 S protein binding to ACE2 of the host cells in vitro.

### Materials and Methods

### Cells and reagents

Recombinant human ACE2 (hACE2) with the Fc region of mouse IgG1 at the C-terminus (hACE2-mFc), SARS-CoV-2 S with poly-His tag at the C-terminus, including full-length S, S1, S2, and RBD, were purchased from Sino Biological (Wayne, PA, USA). HEK293T-hACE2 (NR-52511, BEI Resources, NIAID, NIH), A549-hACE2 (NR-53522, BEI Resources, NIAID, NIH), Hela-hACE2-11 cells (Luo, unpublished) and Vero E6 (CRL-1586, ATCC, Manassas, VA, USA) were maintained in DMEM (GIBCO, NY, USA) supplemented with 10% FBS (GIBCO), penicillin (100 IU/mL), streptomycin (100 μg/mL). BHK-21-/WI-2 cells (EH1011, Kerafast, Boston, MA, USA) were grown in DMEM supplemented with 5% FBS. Calu-3 cells (HTB-5, ATCC, Manassas, VA, USA) were maintained in Collagen type I (Corning) coated plate or flask in DMEM supplemented with 5% FBS. HCT-8 cells (CCL-244, ATCC, Manassas, VA, USA) were cultured in RPMI (GIBCO, NY, USA) supplemented with 10% horse serum (GIBCO). All the cells were cultured in a 5% CO_2_ environment at 37°C and passaged every 2-3 days. The green tea catechins with a purity of >95% (EGCG, ECG, EGC and EC) were purchased from Sigma-Aldrich (St. Louis, MO, USA), and prepared for the stock solution at 10 mM in endotoxin-free water. *Firefly* luciferase substrate was from Promega (Madison, WI, USA).

### Viruses propagation

SARS-CoV-2 (isolate USA-WA1/2020, NR-52281) and human coronavirus (HCoV OC43, NR-52725) were deposited by the Centers for Disease Control and Prevention and obtained through BEI Resources (NIAID, NIH). SARS-CoV-2 viral stocks were generated by infecting Vero E6 cells at ∼90% confluency with SARS-CoV-2 at an MOI of 0.1. At 72 h after infection, supernatants were collected, pooled, and centrifuged at 450 g for 5 min. The resulting stock was aliquoted, tittered using a plaque assay and stored at − 80 °C for future use. All work with live SARS-CoV-2 was performed inside a certified Class II Biosafety Cabinet in the Animal Biological Safety Level-3 (ABSL-3) facility at the University of Nebraska-Lincoln in compliance with all federal guidelines and with the approval of the University of Nebraska-Lincoln Institutional Biosafety Committee. HCoV OC43 was propagated in HCT-8 cells by infection at an MOI of 0.1. HCT-8 cells were incubated at 33°C and 5% CO2 with slow rocking for 6-9 days and then the supernatant was harvested, aliquoted, tittered and stored at − 80°C for further use.

### Cell viability assay

The cytotoxic effects of GTB and its catechins on the cells (HEK293T-hACE2, A549-hACE2, Vero E6 and Huh7) were evaluated by the MTS assay according to the manufacturer’s instruction (Promega, Madison, WI, USA). The cells cultured in 96-well plates were treated with different concentrations (3.125, 6.25, 12.5, 25, 50, and 100 μM) of EGCG or other catechins for 72 h. The cells were then incubated with CellTiter 96® AQueous One Solution Reagent containing MTS and phenazine methosulfate at 37°C for 4 h in darkness. Absorbance at 490 nm was measured by a plate reader (Spectra Max i3, Molecular Devices, Sunnyvale, CA, USA). Cell viability was calculated and normalized with untreated control, which was defined as 100%.

### Plasmids and pseudoviruses

For generating the pseudovirus bearing SARS-CoV-2 S, the vector pCAG-SARS-CoV-2-Sd18 encoding human codon-optimized wildtype (WT) S gene of SARS-CoV-2 with C-terminal 18 aa deletion (Sd18) was constructed using NEBuilder® HiFi DNA Assembly cloning kit (NEB, E5520S). To enhance the expression of Sd18 protein for better packaging efficiency of pseudovirus, the Sd18 expression cassette from the CMV-driven vector pcDNA3.1-SARS2-S, a gift from Fang Li (Addgene Cat # 145032), was transferred to a CAG-driven vector pCAG-Flag-SARS-CoV-2-S (gift from Peihui Wang, Shandong University) via *EcoR*I/*Not*I sites and PCR with forward primer (5’-TGGTGGAATTCTGCAGATgctagcatgtt-tgtcttcctg-3’) and reverse primer 5’-(CCTGCACCTGAGGAGTTActtgcagcagctgccgcagga-3’). PCR was performed with `Phusion High-Fidelity PCR Master Mix` kit (ThermoFisher Scientific, F531S) and purified using the Monarch PCR & DNA Cleanup Kit (NEB, T1030S). After verification with Sanger sequencing, this pCAG-SARS-CoV-2-Sd18 vector (WT-Sd18) was used as the backbone for the subsequent cloning of single and full-set mutations S of variants. For the introducing 1-4 mutations in S variant (e.g. CA) or shared multiplexed mutants (e.g. E484K, N501Y, D614G), one step of NEBuilder HiFi cloning was performed using WT-Sd18 as the backbone vector with *EcoR*I/*PpuM*I double digestion and 1-4 fragments of PCR products with mutagenic primers targeting each mutation site (**Table S1**). For the introducing 5-9 mutations in S variant (e.g. UK, SA), two steps of NEBuilder HiFi cloning were performed by generating N-terminal mutant with *EcoR*I/*PpuM*I and C-terminal mutant with *PpuM*I/*Not*I with 1-5 fragments of PCR products, and then either N-terminal or C-terminal mutant as the backbone vector for further cloning of other mutant regions. The pCAG-hACE2-Flag plasmid was a gift from Peihui Wang (Shandong University). The nucleotide sequences of all the plasmids were confirmed by Sanger sequence analysis (Genewiz).

The vesicular stomatitis virus (VSV)-based pseudovirus bearing S has been extensively used for SARS-CoV-2 studies^[33-36]^. The viral backbone was provided by VSVΔG that packages reporter expression cassettes for *firefly* luciferase (VSVΔG-Luc) or GFP (VSVΔG-GFP) instead of VSVG in the genome. Briefly, BHK-21/WI-2 cells were transfected with S encoding plasmid (15 μg for 10 cm dish) using PEIpro transfection reagent (Polyplus, Illkirch, France) following the manufacturer’s instruction. At 24 h post transfection, cells were infected with VSVΔG-based reporter pseudovirus bearing VSVG (VSVG-ΔG-reporter, Kerafast, Cat# EH1020-PM and Cat# EH1019-PM) at MOI of 4 for 2 h, then cells were washed with PBS three times, and cultured in new complete medium. At 24 h post infection, the pseudovirus-containing culture supernatant was harvested and centrifuged at 500 g for 5 min to remove cell debris. The cell-free supernatant was then filtered through a 0.45 μm polyethersulfone membrane and stored at -80°C in 100 μL aliquots for future use.

To study SARS-CoV-2 entry into target cells, the containing lentivirus-based S pseudovirus bearing β-lactamase-Vpr (BlaM-Vpr) chimeric protein was generated as previously described[37]. Briefly, HEK293T cells were co-transfected with SARS-CoV-2 S (10 μg), psPax2 (10 μg, a gift from Didier Trono, Addgene Cat# 12260) and HIV-1 YU2 Vpr β-lactamase expression vector pMM310 (10 μg, NIH AIDS Reagents Cat #11444) using PEIpro transfection reagent (Polyplus, Illkirch, France). The medium was changed at 4 h post-transfection, then total supernatant was collected at 48 h post-transfection and centrifuged at 500 g for 5 min to remove cell debris, and filtered through a 0.45 μm polyethersulfone membrane. The viruses were then concentrated (1:100) by centrifugal ultrafiltration with Amicon® Ultra-15 filter device (50,000 NMWL, Millipore, Burlington, MA, USA). Virus stocks were stored in a single-use aliquot and used only once to avoid inconsistencies due to repeated freezing-thawing cycles.

### Green tea beverage (GTB) preparation

Matcha green tea bags were purchased from Ito En (Shibuya City, Tokyo, Japan). GTB was prepared as instructed by the manufacturer. Briefly, the tea bag was soaked in 150 mL of hot (95°C) water in a teacup for 30 sec and then shake 5-6 times before being discarded. GTB was centrifuged at 4,000 rpm for 10 min to remove undissolved particles, and the supernatant was then filtrated through a 0.22 μm filter. The filtered GTB was aliquoted and stored at -80°C.

### Green tea and its catechin treatment

Cells or the viruses were preincubated with or without GTB or its catechins at different concentrations prior to the infection. For the VSVΔG-Luci based S pseudovirus detection, the culture supernatant was aspirated gently, and the cells were rinsed with 1× PBS, and lysed with 1× Passive Lysis Buffer (Promega, Madison, WI, USA). The cell lysate was transferred to a 96-well white plate, and 100 μL of *firefly* luciferase substrate (Promega, Madison, WI, USA) was added to each well for the detection of luminescence using a microplate luminometer (Infinite 200 PRO, Tecan). For VSVΔG-GFP based pseudovirus detection, GFP fluorescent images were captured under Nikon A1R confocal microscopy.

To confirm the broad-spectrum antiviral effect of EGCG, the cells (HEK293T-hACE2 and HCT-8) were pretreated with EGCG (100 μM) or other catechins for 30 min prior to incubation with live SARS-CoV-2 (USA isolate, 2019-nCoV/USA-WA1/2020) or live HCoV OC43. The cells were then washed to remove the free viruses 1 h (live SARS-CoV-2) or 2 h (live HCoV OC43) post-inoculation and cultured in complete medium containing the GTB or catechins. Supernatant from live SARS-CoV-2-infected HEK293T-hACE2 cells was collected at 48 h post-infection for RNA extraction. HCT-8 cells infected with HCoV OC43 were collected at day 4 post-infection for RNA extraction.

### Pseudovirus entry assay

The BlaM-Vpr-containing lentivirus-based pseudovirus bearing SARS-CoV-2 S was used to investigate the effect of EGCG on cellular entry of SARS-CoV-2. After the virus entry into the host cells, BlaM is released into the cytoplasm of the infected cells. Virus fusion and entry can be detected by enzymatic cleavage of CCF4-AM, a fluorescent substrate of BlaM, delivered into the host cell cytoplasm. The β-lactam ring in CCF4 is cleaved by BlaM, which changes the emission spectrum of CCF4 from 520 nm (green) to ∼450 nm (blue), allowing the entry of virus to be quantified using flow cytometry. BlaM-Vpr-containing lentivirus-based S pseudovirus was preincubated with EGCG for 30 min at 37°C before being incubated with HEK293T-hACE2 cells for 4 h. CCF4-AM containing solution was then added to cell culture and incubated for 2 h at 22°C to prevent further fusion and to allow substrate cleavage by BlaM. Percentage of CCF4 cleavage was assessed by flow cytometry on a FACSCanto II (BD Bioscience) and analyzed using FlowJo software (Tree Star Inc., Ashland, OR). Non-infected cells treated with CCF4-AM were used to set the gate for uncleaved CCF4, which was set to discriminate entry and non-entry at a tolerance of <1% false positives.

### Immunofluorescence and confocal microscopy

A549 cells were pre-seeded in 8-well chambered coverslip (C8-1.5H-N, Cellvis, Mountain View, CA, USA), and transfected with the plasmid expressing hACE2 gene. Twenty-four hours later, the cells were incubated with recombinant SARS-CoV-2 S with His-tag in the presence or absence of EGCG for 6 h, and then washed three times with PBS to remove unbound protein, fixed with 4% paraformaldehyde in PBS for 10 min. The cells were subsequently blocked in PBS containing 1% BSA for 1 h, and then incubated with goat anti-ACE2 (R&D system, 1:100) and mouse anti-His-tag (Proteintech, 1:400 dilution) overnight at 4°C, followed by staining with donkey anti-goat IgG antibody conjugated with Alexa Fluor 594 (Invitrogen, 1:1000) and donkey anti-mouse IgG antibody conjugated with Alexa Fluor 488 (Invitrogen, 1:1000) for 60 min at room temperature in the dark. Nuclei were stained with 1 μg/mL Hoechst (Invitrogen). All the images were acquired using confocal microscopy (Nikon A1R, Nikon, Japan).

### Western blot

To determine the incorporation of the S in VSVΔG-based pseudoviruses, supernatant (10 mL) was collected from BHK-21/WI-2 cell cultures and concentrated through a 10% sucrose cushion by centrifugation at 8,000× g for 3 h. The layers of supernatant and sucrose were removed, and the resulting viral pellets were harvested in RIPA buffer (100 μL). The prepared pseudoviruses (65 μL) were mixed with 4× LDS sample buffer (25 μL) and 10× sample reducing agent (10 μL). The mixture was denatured for 10 min at 70°C. The sample (20 μL) was subjected to western blot assay and S incorporated on the pseudovirus surface was detected by convalescent plasma (1:500 dilution) from COVID-19 patient and goat anti-human IgG (1:1000 dilution) (ImmunoReagents, Raleigh, NC, USA).

Cells were harvested in RIPA buffer supplemented with the protease and phosphatase inhibitor mixture. The protein concentration was determined by the bicinchoninic acid (BCA) protein assay (ThermoScientific, Rockford, IL, USA) Proteins were separated in Bis-Tris gradient gel and analyzed by antibody using the following antibodies and dilutions: ACE2 (MA5-31395, Invitrogen, 1:1000), GAPDH (2118S, Cell Signaling Technology, 1:1000), His-tag (66005-1-Ig, Proteintech, 1:5000), SARS-CoV-2 Nucleocapsid (MA5-35943, Invitrogen, 1:2000), and hamster GAPDH (10494-1-AP, Proteintech, 1:5000). Secondary antibodies were anti-rabbit (7074S, Cell Signaling Technology) or anti-mouse (7076S, Cell Signaling Technology) IgG conjugated with horseradish peroxidase. Protein bands were visualized with enhanced ECL substrate and Invitrogen iBright FL1500 Imaging System.

## ELISA

Recombinant hACE2 protein (2 μg/mL) was precoated onto an ELISA plate at 4°C overnight. The coated plate was blocked with 2% BSA in PBST (pH=7.4) at room temperature for 1 h, and incubated with PBS with or without EGCG at different concentrations for 30 min. The serially diluted SARS-CoV-2 S with His tag was added to the plate. HRP-conjugated anti-His tag antibody (HRP-66005, Proteintech, 0.2 μg/mL) was sequentially added as a detection antibody. Peroxidase substrate solution (TMB) and 1M H_2_SO4 stop solution were used and the absorbance (OD 450 nm and 570 nm) was read by a microplate reader (Spectra Max i3, Molecular Devices, Sunnyvale, CA, USA).

### RNA Extraction and Reverse Transcription Quantitative PCR (RT-qPCR)

Viral RNA was extracted from the cell supernatant using QIAamp Viral RNA mini kit (Qiagen, Cat# 52906) as recommended by the manufacturer and quantified using C1000 Thermal Cycler and the CFX96 Real-time system (Bio-Rad). Standards were prepared by serial dilution of 2019-nCoV_N_Positive Control (IDT, Cat# 10006625) at different concentration and Control RNA from heat-inactivated SARS-CoV-2 (BEI Resources, Cat# NR-52347) was used as RT-qPCR positive control. For 20 µL of RT-qPCR reaction, 5 µL of extracted viral RNA, TaqMan Fast Virus 1-Step master mix (ThermoFisher Scientific, Cat# 4444434) and 2019-nCov CDC EUA kit (IDT, Cat# 10006770) for N1 primer-probe combination was used: 2019-nCoV_N1-F (GACCCCAAA ATCAGCGAAAT), 2019-nCoV_N1-R (TCTGGTTACTGCCAGTT-GAATCTG), 2019-nCoV_N1-P (FAM-ACCCCGCATTACGTTTGGTGGACC-BHQ1). HCoV OC43 RNA copies were measured by the RT-qPCR for the viral M gene expression with the forward primer (5’-ATGT-TAGGCCGATAATTGAGGACTAT-3’) and reverse primer (5’-AATGTAAAGATGGCCGCGTATT-3’) using QuantStudio 3 system (Applied Biosystems, ThermoFisher Scientific) with Powerup SYBR Green master mix (Thermo Fisher Scientific, Cat# A25743). Intracellular level of viral RNA was normalized with GAPDH using forward primer (5′-GGTGGTCTCCTCTGACTTCAACA-3′) and reverse primer (5′-GTTGCTGTAGC-CAAATTCGTTGT-3′).

### Statistical Analysis

All data were presented as mean ± SD. Data were analyzed statistically using unpaired *t* tests when two groups were being compared or by one-way ANOVA for multiple comparisons using GraphPad Prism 8.0.1. Nonlinear regression (four parameters) was performed to calculate IC_50_ values using the inhibitor versus response and EC_50_ values using agonist versus response least-squares fit algorithm. For all data, significance was presented as \**P* < 0.05, ** *P* < 0.01 and *** *P* < 0.001.

## Results

### Generation of VSVΔG-based pseudoviruses bearing SARS-CoV-2 Spikes (WT and variants)

Pseudoviruses have been extensively used in diagnostics, vaccines, and high-throughput screening of entry inhibitors for some BSL-3/BSL-4 level pathogens^[34, 35, 38, 39]^. VSVΔG-based pseudovirus is an ideal model for *in vitro* studies of SARS-CoV-2 because the native envelope protein of VSV can be replaced with S of SARS-CoV-2 **(Supplemental Fig. 1**). Therefore, we generated several VSVΔG-based pseudoviruses bearing S with single mutation (K417N, E484K, N501Y, D614G) or S with full-set mutations of the newly emerged variants (UK-B.1.1.7, SA-B.1.351, and CA-B.1.429) (**Fig. 1A and 1B**). To confirm the S incorporation into VSVΔG-based pseudoviruses, we examined the S expression in the viruses and the producer cells by western blot using convalescent plasma from COVID-19 patients. As shown in **Supplemental Fig. 2A**, specific bands could be detected in both the pseudovirions and the cell lysates of the producer cells. In contrast, no specific bands were observed in the control. We next examined whether the pseudoviruses bearing SARS-CoV-2 S can infect HEK293T-hACE2 cells. As shown in **Supplemental Fig. 2B**, the pseudoviruses bearing WT S or D614G S could infect the hACE2-expressing cells evidenced by increased luciferase activity after infection. To exclude the possibility of infection was deriving from the residual trace VSVG-pseudovirus during packaging of the S-pseudoviruses, we performed the neutralization assay with the specific antibody against VSVG. While the antibody completely blocked VSVG-pseudovirus infection, it had little effect on the infectivity of pseudoviruses bearing either WT or D614G S (**Supplemental Fig. 2C**). We next examined the specificity of S-pseudoviruses to infect the hACE2-expressing cells using convalescent plasma neutralization assay.

**Figure 1.**
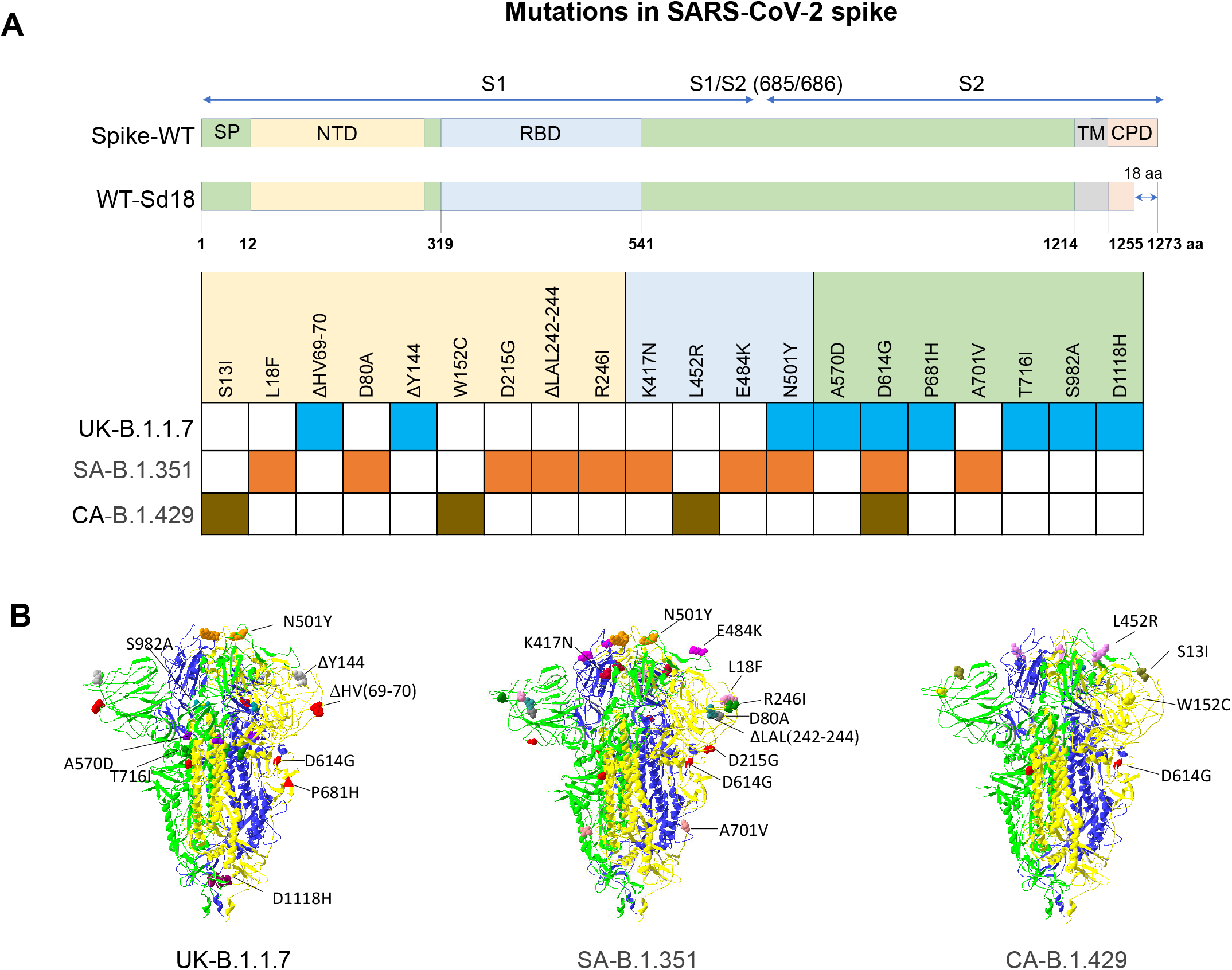
Schematic overview of the S proteins from the SARS-CoV-2 variants in this study. (**A**) The indicated domains and elements, including signal peptide (SP), N-terminal domain (NTD), receptor binding domain (RBD), transmembrane domain (TM) and cytoplasmic domain (CPD), are marked. All the mutations found in the UK variant (UK-B.1.1.7), South Africa variant (SA-B.1.351), and California variant (CA-B.1.429) were shown. All the spike, including wildtype and single or full-set mutations of variants were constructed to generate SARS-CoV-2 pseudovirus. To enhance the packaging efficiency, 18 amino acids at C terminal of spike were depleted. (**B**) The location of the mutations in the context of the trimer spike protein domain. 3D trimer structure of SARS-CoV-2 S protein at closed state (PDB ID: 7DDD) was generated using SwissPdb viewer. Three SARS-CoV-2 S monomers are shown as ribbon diagram in green, yellow and blue colors. The mutated residues are shown as space filling spheres in different colors to distinguish the different mutations. The mutations are labelled only in one of the monomers. Red triangle in UK variant depicts approximate location of P681H, which was not included in the model.

**Figure 2.**
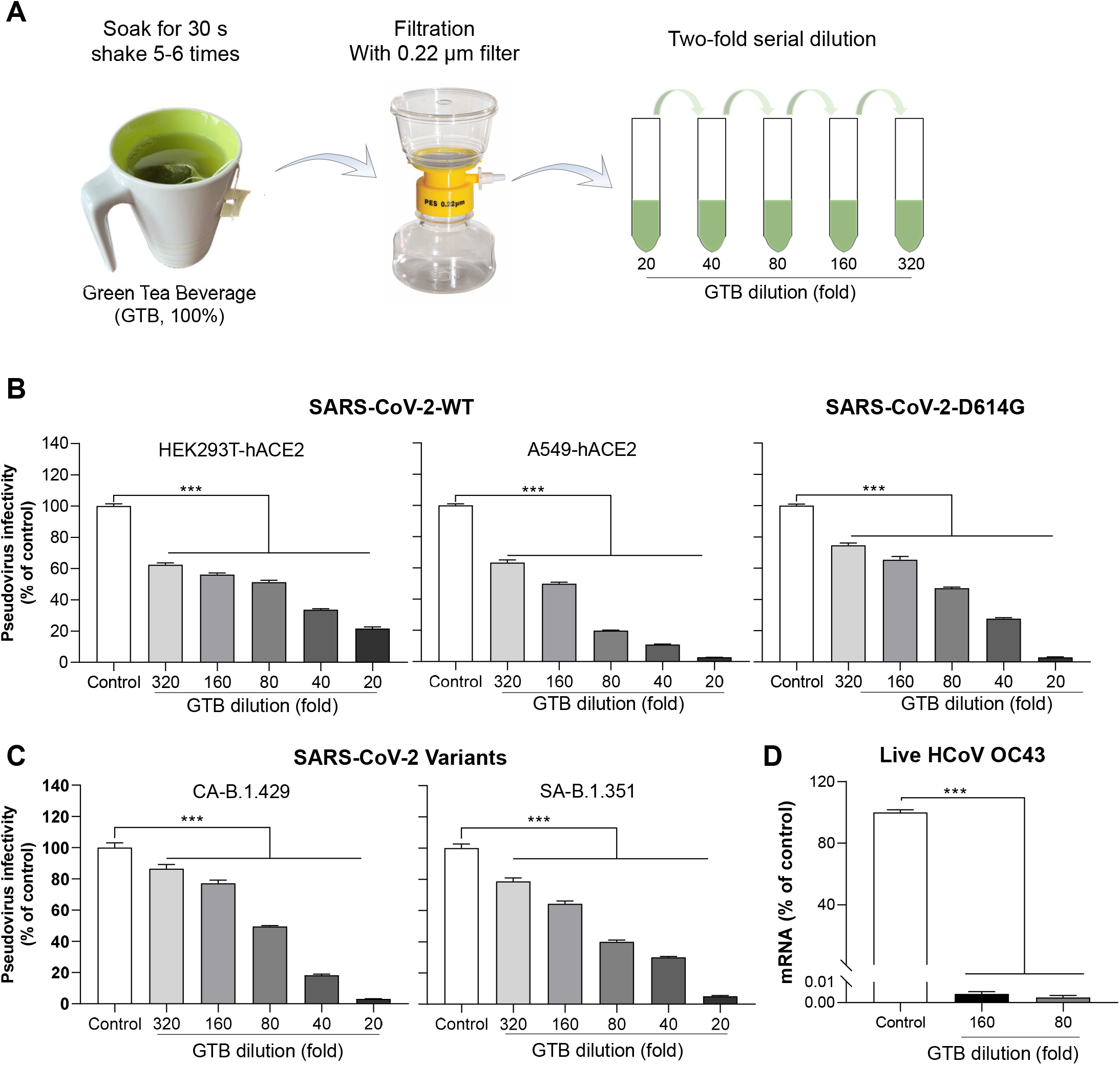
Green tea beverage (GTB) inhibits SARS-CoV-2 pseudovirus and variants infection. (**A**) GTB was prepared by soaking 1.8 g of green tea bag in 150 mL water at 90-95°C for 30 sec. After centrifugation at 4,000 rpm for 10 min, supernatant was collected and filtrated through 0.22 μm filter, which was defined as 100% GTB. A 2-fold serial dilution of GTB was prepared as indicated. (**B**) The VSVΔG-based pseudovirus bearing SARS-CoV-2 S (SARS-CoV-2-WT, D614G) was pre-incubated with diluted GTB for 30 min prior to infection of HEK293T-hACE2 or A549-hACE2 cells. At 48 h post-infection, cells were lysed and luciferase activity was measured to assess the viral infectivity expressed as a percentage relative to that of the control (untreated). (**C**) The VSVΔG-based pseudovirus bearing full-set mutant SARS-CoV-2 S (CA-B.1.429, SA-B.1.351) was pre-incubated with diluted GTB for 30 min prior to infection of HEK293T-hACE2 cells. At 48 h post-infection, cells were lysed and luciferase activity was measured to assess the viral infectivity expressed as a percentage relative to that of the control (untreated). (**D**) HCoV OC43 was pre-incubated with diluted GTB for 30 min prior to infection of HCT-8 cells. Intracellular HCoV OC43 RNA were determined at day 4 post-infection by RT-qPCR. Data are shown as mean ± SD, representative of two independent experiments with 3 replicates.

The convalescent plasma from the COVID-19 patients could specifically block the infection of the pseudoviruses bearing either WT or D614G S as evidenced by the typical four-parameter inhibition curves (**Supplemental Fig. 2D**).

### Green tea beverage (GTB) and its catechins are non-toxic to cells

To study the antiviral effect of green tea beverage (GTB), we prepared GTB based on the manufacturer’s instruction as illustrated in **Fig. 2A**. Before evaluating the antiviral activity of GTB and the green tea catechins (EGCG, ECG, EGC and EC), we examined whether they affect the viability of four cell types (HEK293T-hACE2, A549-hACE2, and HCT-8) used in this study. As shown in **Supplemental Fig. 3A**, GTB had little cytotoxicity to the cells at the dilutions used in the experiments. In addition, the green tea catechins treatment at the concentration as high as 100 μM for as long as 72 h had little cytotoxic effect on the cells (**Supplemental Fig. 3B**).

**Figure 3.**
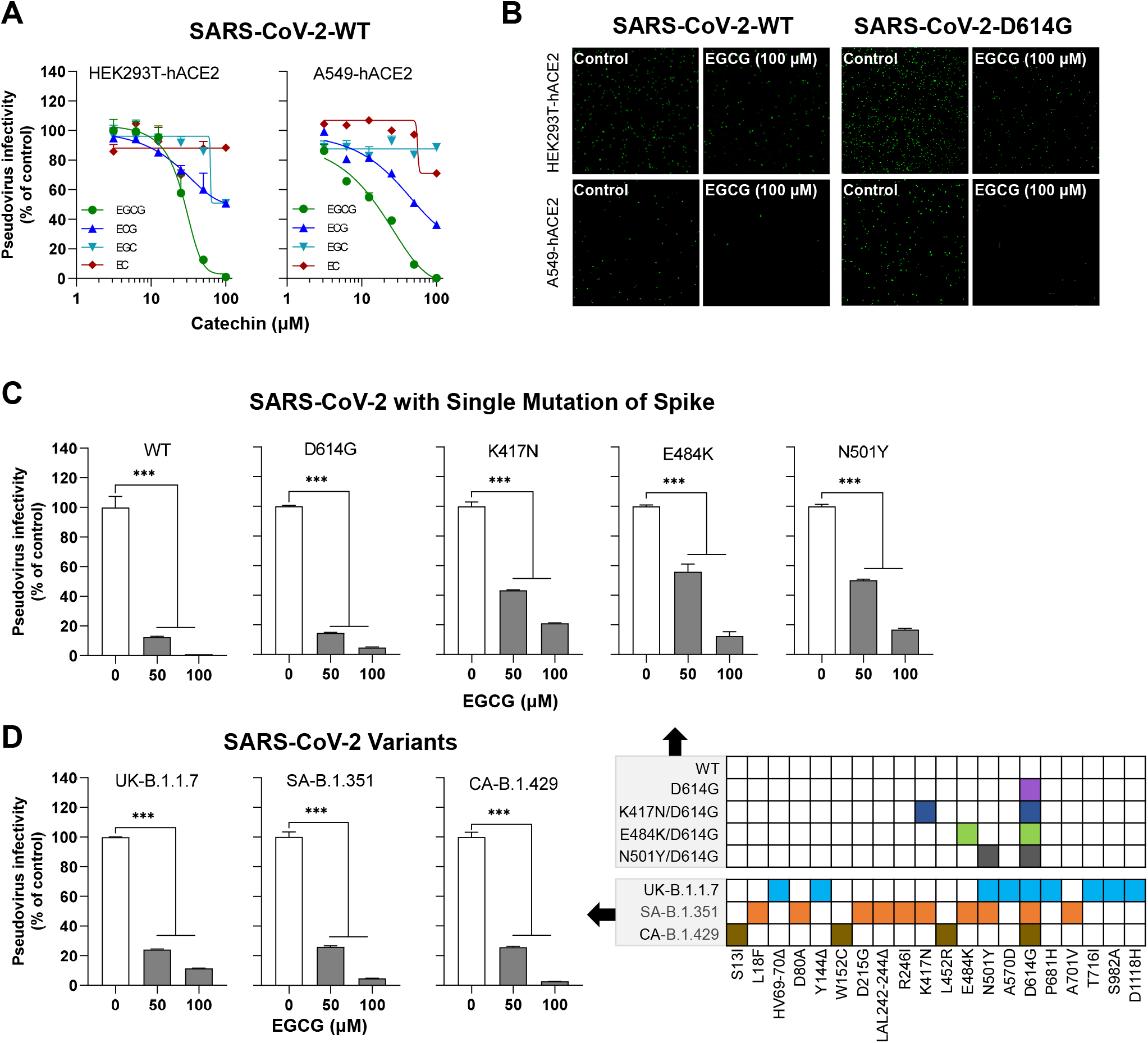
EGCG inhibits SARS-CoV-2 spike (WT and variants)-mediated infection. (**A**) HEK293T-hACE2, A549-hACE2 and VeroE6 cells were treated with the indicated doses of green tea catechins (EGCG, ECG, EGC and EC) for 30 min prior to infection with the pseudovirus bearing SARS-CoV-2 S. (**B**) HEK293T-hACE2 cells and A549-hACE2 cells were pretreated with indicated doses of EGCG for 30 min prior to infection of the pseudovirus bearing SARS-CoV-2 S WT or D614G. GFP fluorescent images were captured by Nikon A1R confocal microscope. (**C**) HEK293T-hACE2 cells were incubated with the indicated doses of EGCG for 30 min prior to infection of the pseudovirus bearing SARS-CoV-2 spike with mutations indicated. (**D**) HEK293T-hACE2 cells were pretreated with EGCG for 30 min prior to infection of the pseudovirus bearing SARS-CoV-2 S of four newly emerged variants. Cells were lysed and luciferase activity was measured at 48 h post-infection. Viral infectivity was assessed by luciferase activity, which is expressed as a percentage relative to that of the control (untreated). Data are shown as mean ± SD, representative of two independent experiments. \*\*\**P*<0.001.

### GTB inhibits infection of SARS-CoV-2 pseudovirus and live human coronavirus

GTB preincubation with the pseudovirus bearing SARS-CoV-2 S (WT or D614G) dose-dependently inhibited the viral infectivity (**Fig. 2B**). In addition, GTB preincubation significantly suppressed the infectivity of the pseudoviruses bearing S with the full-set mutations of newly emerged SARS-CoV-2 variants (CA-B.1.429, and SA-B.1.351) in a dose-dependent manner (**Fig. 2C**). Furthermore, we examined whether the GTB can inhibit live HCoV OC43 nfection. As shown in **Fig. 2D**, GTB at dilution as much as 160-fold could completely block HCoV OC43 infection of HCT-8 cells.

### EGCG inhibits SARS-CoV-2 spike (WT and variants)-mediated infection

To determine the component(s) in green tea responsible for the viral inhibition, we examined anti-SARS-CoV-2 activity of the major active ingredients (catechins) in green tea, which include EGCG, EGC, ECG and EC. EGCG accounts for more than 50% of the total amount of catechins in green tea^[40]^. As demonstrated in **Fig. 3A**, three (EGCG, EGC and ECG) out of 4 catechins could dose-dependently inhibit SARS-CoV-2 WT S-mediated pseudovirus infection of HEK293T-hACE2 and A549-hACE2 cells (**Fig. 3A**). Among three green tea catechins, EGCG was the most potent inhibitor of the viral infection (**Fig. 3A**). Therefore, we focused on EGCG for the subsequent experiments.

To visualize the anti-SARS-CoV-2 activity of EGCG in the host cells, we examined the inhibitory effect of EGCG on VSVΔG-GFP-based pseudoviruses bearing WT S or D614G S. As shown in **Fig. 3B**, there was a significant reduction of the GFP positive cells in both HEK293-hACE2 and A549-hACE2 cells treated with EGCG. In addition to its inhibitory effect on SARS-CoV-2 WT S, EGCG also efficiently inhibited infectivity of the pseudoviruses bearing WT S and S with single mutation (D614G, K417N, E484K and N501Y) (**Fig. 3C**). These mutations occur in the RBD of the newly emerged SARS-CoV-2 variants, which are critical for the viral infection and evasion against neutralizing antibody^[41-44]^. Importantly, EGCG significantly suppressed the infectivity of the pseudoviruses bearing full-set mutations S of the newly emerged variants (UK-B.1.17, SA-B.1.351, and CA-B.1.429) (**Fig. 3D**).

To rule out the possibility that EGCG present in the cell lysates can influence luciferase activity analysis, we added EGCG to the cell lysates before the luciferase activity assay. We found no effect of EGCG on luciferase activity (**Supplemental Fig. 4A**). In addition, EGCG did not affect luciferase gene expression in the cells transfected with the plasmid with luciferase gene (**Supplemental Fig. 4B**).

**Figure 4.**
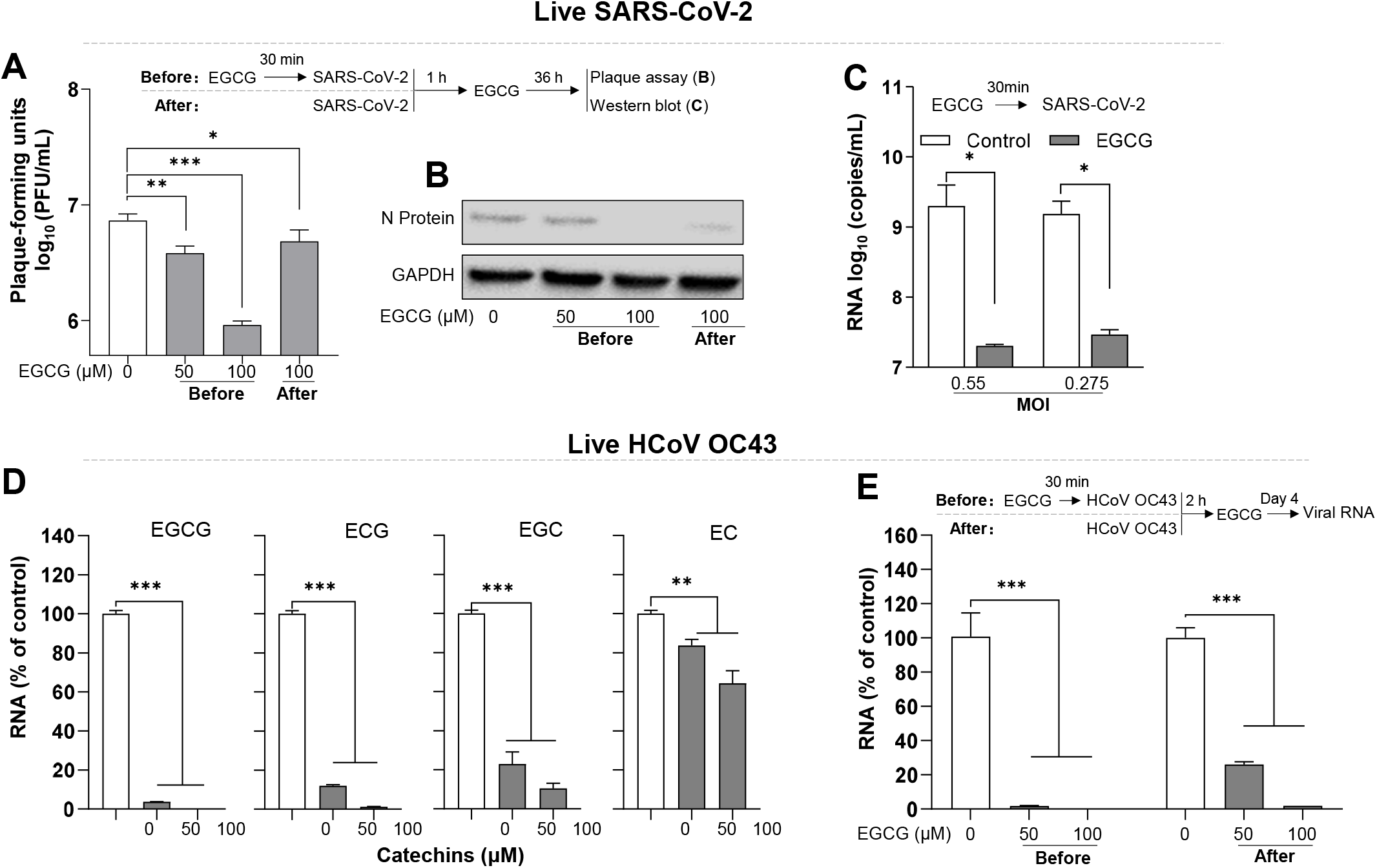
EGCG inhibits live SARS-CoV-2 and HCoV OC43 infection. (**A**) Human lung epithelial cells (Calu-3) were treated with EGCG (50,100 μM) before or after live SARS-CoV-2 (USA isolate) infection at MOI=5. Cells were washed 3 times with pre-warmed medium to remove free virus at 1 h post-infection, and then maintained in complete medium containing EGCG for 36 h. Supernatants were collected and plaque assay was carried out using Hela/ACE2-11 cells. (**B**) Cells from (**A**) were lysed and subjected to western blot to detect SARS-CoV-2 N protein. (**C**) HEK293T-hACE2 cells were pretreated with EGCG (100 μM) for 30 min prior to live SARS-CoV-2 (USA isolate) infection at MOI=0.055 or 0.275. After 1 h virus adsorption in the presence of EGCG, the cells were washed twice with pre-warmed medium, and then maintained in complete medium containing EGCG for 48 h. SARS-CoV-2 RNA copies from culture supernatant were determined at 48 h post-infection by RT-qPCR. (**D**) HCT-8 cells (susceptible cells for HCoV OC43) were pretreated with catechins at indicated dose for 30 min prior to HCoV OC43 infection at MOI=0.1. After 2 h virus adsorption, the cells were washed twice with pre-warmed medium, and then maintain in complete medium containing EGCG. (**E**) HCT-8 cells were treated with EGCG (50,100 μM) before or after live HCoV OC43 infection at MOI=0.1. After 2 h virus adsorption, the cells were washed twice with pre-warmed medium, and then maintain in complete medium containing EGCG. Intracellular HCoV OC43 RNA were determined at day 4 post-infection by qRT-PCR. Data are shown as mean ± SD, representative of two independent experiments with 3 replicates. \**P*<0.05, \*\**P*<0.01, \*\*\**P*<0.001.

### EGCG inhibits live SARS-CoV-2 and HCoV OC43 infection

To further determine the significance of anti-SARS-CoV-2 activity of EGCG, we examined the effect of EGCG on live SARS-CoV-2 infection in human lung epithelial cell (Calu-3). EGCG could efficiently suppress live SARS-CoV-2 infection (**Fig. 4A**). While the inhibitory effect on the virus was observed in the cells treated with EGCG either before or after infection, the pretreatment of the cells with EGCG exhibited higher inhibitory activity than its treatment after infection (**Fig. 4A and 4B**). We next confirm the results using HEK293T-hACE2 and found that EGCG pretreatment of the cells potently (>95%) inhibited live SARS-CoV-2 infection with different MOI (**Fig. 4C**).

We also studied effects of EGCG and other three catechins on live human coronavirus (HCoV OC43) infection. As shown in **Fig. 4D**, among four green tea catechins, while EC was the least effective in suppressing the virus, EGCG was the most potent in the viral inhibition (**Fig. 4D**). Similar to its effect on SARS-CoV-2, EGCG exhibited the antiviral activity of HCoV OC43 in the cells treated either before or after infection, and the pre-treatment was more efficient than the post-treatment (**Fig. 4E**).

### EGCG blocks SARS-CoV-2 S-mediated viral entry

To further determine the anti-SARS-CoV-2 mechanism of EGCG, we examined the ability of EGCG to block the virus entry under four different experimental conditions (**Fig. 5A**): EGCG was premixed with the pseudovirus or EGCG was added the cell cultures either before, during or after the viral infection. As demonstrated in **Fig. 5A**, although EGCG treatment after the infection had little effect, the significant viral inhibition was observed when the cells were treated with EGCG before or during the viral infection. The highest viral inhibition occurred when EGCG was premixed with the virus prior to the infection (**Fig. 5A**). To further determine whether EGCG can block SARS-CoV-2 entry, we performed BlaM-based virus entry assay using lentivirus-based pseudovirus containing BlaM-Vpr chimeric protein and bearing SARS-CoV-2 S. As shown in **Fig 5B**, EGCG pretreatment significantly blocked the virus entry into the target cells as evidenced by reduced CCF4 cleavage.

**Figure 5.**
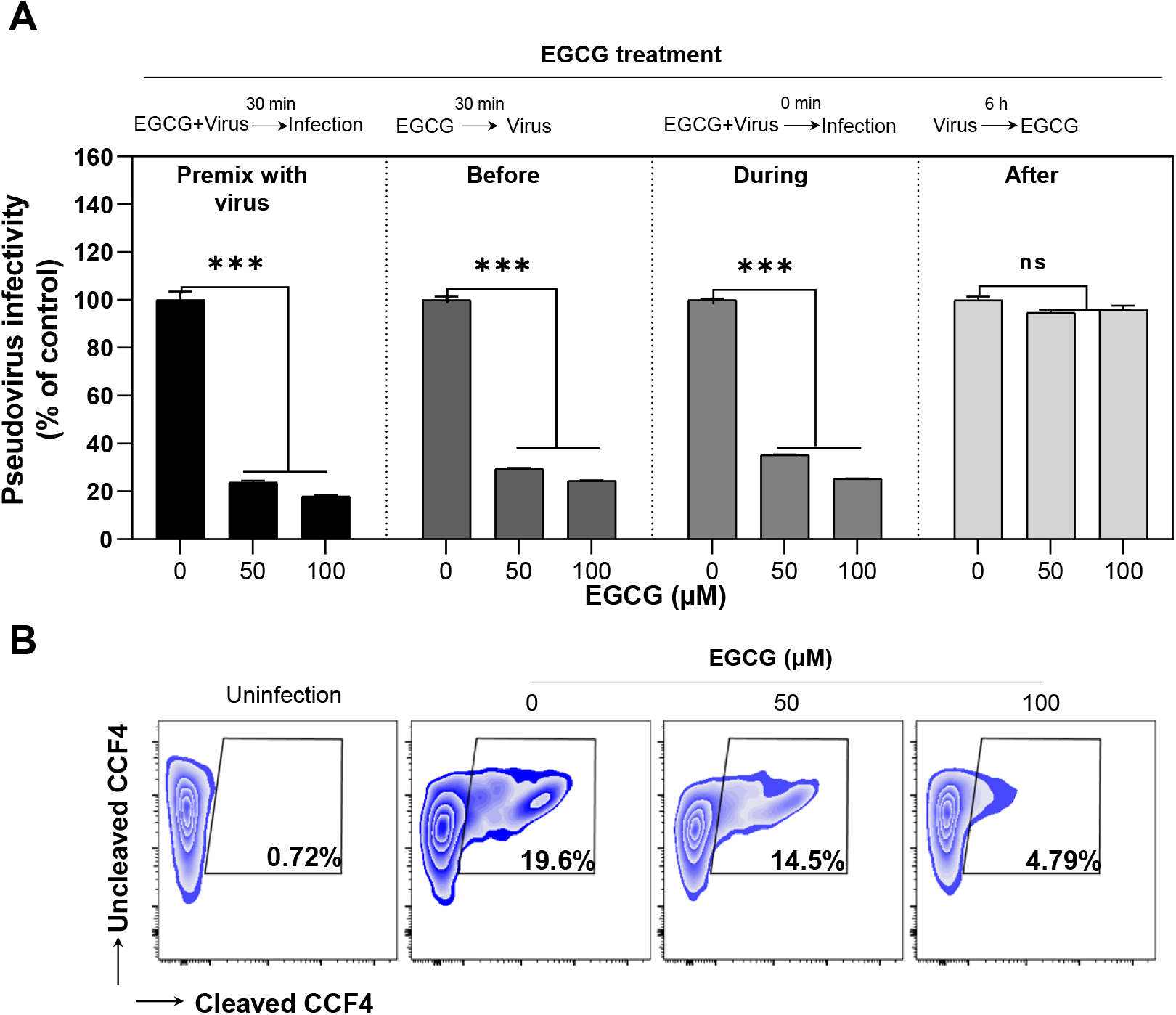
EGCG inhibits SARS-CoV-2 entry. (**A**) HEK293T-hACE2 cells were infected with SARS-CoV-2 pseudovirus and treat with EGCG under 4 different conditions as indicated. Cells were lyzed at 48 h postinfection and luciferase activity was measured. Data are shown as mean ± SD of three replicates. (**B**) BlaM-containing SARS-CoV-2 S pseudotyped lentivirus was pre-incubated with EGCG for 30 min at 37°C, subsequently infected HEK293T-hACE2 cells for 4 h to allow virus entry. The cells were then loaded with CCF4-AM to monitor cleavage and shift in fluorescence output for evidence of S mediated viral entry into cells. Percentage of CCF4 cleavage was assessed by flow cytometry on a FACSCanto II. Non-infected cells treated with CCF4-AM were used to set the gate for uncleaved CCF4, which was set to discriminate entry.

### EGCG inhibits SARS-CoV-2 S binding to ACE2

The initial and key step for SARS-CoV-2 infection is binding to the ACE2 receptor of host cells via the viral S^[45]^. To study the specific effect of EGCG on blocking SARS-CoV-2 entry, we examined whether EGCG interferes with the engagement of SARS-CoV-2 S and ACE2 receptor. We prepared His-tagged SARS-CoV-2 full-length S to study its binding affinity to ACE2 receptor of the host cells treated with or without EGCG. As shown in **Fig. 6A**, EGCG could block the binding of full-length S to ACE2. To further determine the specific binding site(s) affected by EGCG, we examined the binding affinity of the S subunits (S1, S2 and RBD) to ACE2. As shown in **Fig. 6B**, the S1 subunit had the high binding affinity to ACE2, which could be blocked by EGCG treatment. In contrast, the S2 subunit showed little binding affinity to ACE2 regardless of EGCG treatment (**Fig. 6C**). In addition, EGCG significantly reduced the binding of RBD to ACE2 (**Fig. 6D**).

**Figure 6.**
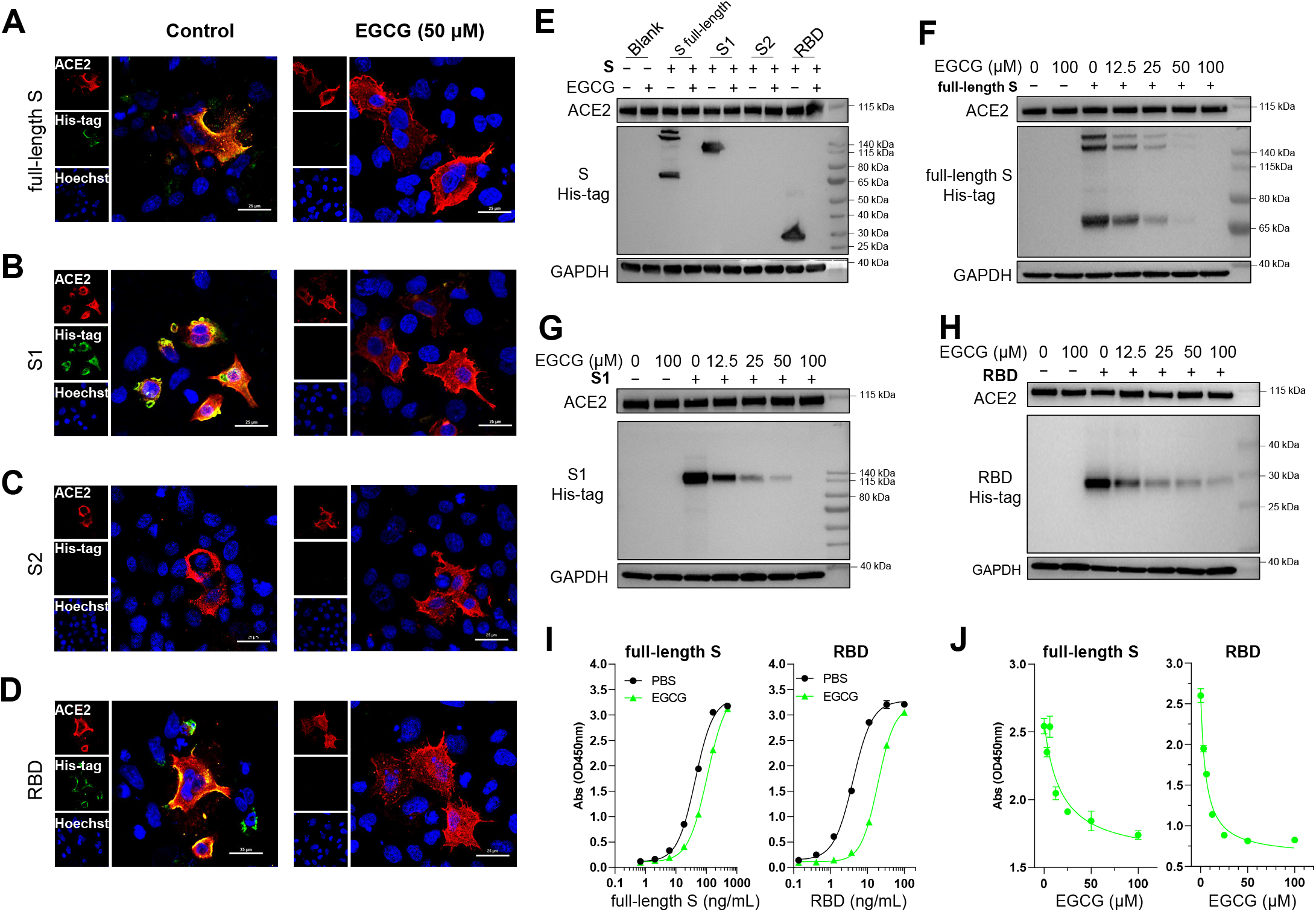
EGCG blocks SARS-CoV-2 S binding to ACE2. (**A-D**) A549 cells were transfected with pCAG-hACE2 plasmid. Twenty-four hours later, the cells were incubated with medium containing His-tagged SARS-CoV-2 S (full-length S, S1, S2 and RBD, 1 μg/mL) with or without EGCG (50 μM) for 6 h, and subsequently incubated with goat anti-ACE2 and mouse anti-His-Tag antibodies, following stained with donkey anti-goat IgG antibody conjugated with Alexa Fluor 594 and donkey anti-mouse IgG antibody conjugated with Alexa Fluor 488. Nuclei were stained with Hoechst. All images were obtained by confocal microscopy (Nikon A1R, Nikon, Japan). The scale bar in each panel indicates 25 μm. (**E**) VeroE6 cells were incubated with medium containing His-tagged SARS-CoV-2 S (full-length S, S1, S2 and RBD, 1 μg/mL) with or without EGCG (50 μM) for 6 h, and then washed 3 times with PBS to remove unbound protein. Cell lysate was subsequently subjected to western blot analysis to detect the binding of His-tagged S to ACE2. GAPDH was set as internal control of western blot. (**F-H**) SARS-CoV-2 S (full-length S, S1 and RBD, 1 μg/mL) was premixed with EGCG at the indicated dose, and the premixture was then incubated with VeroE6 cells for 6 h, free S protein was removed with PBS triple washing. Western blot analysis was performed to assess the binding of S protein to ACE2. (**I**) Immobilized hACE2 protein (Fc tag) was precoated at 2 μg/mL at 4°C overnight, and 3-fold serially-diluted His-tagged SARS-CoV-2 full-length S (500∼0.686 ng/mL) and RBD (100∼0.137 ng/mL) were used to test the binding affinity to ACE2 in presence or absence of EGCG (50 μM) by ELISA assay. (**J**) Immobilized hACE2 protein (Fc tag) was precoated at 2 μg/mL at 4°C overnight, His-tagged full-length S (100 ng/mL) and RBD (10 ng/mL) were premixed with EGCG at indicated dose (0∼100 μM), and then used to test the binding affinity to ACE2 by ELISA assay. The data shown are representative of two independent experiments.

To confirm the inhibitory effect of EGCG on the S binding to ACE2, we performed western blot analysis to determine the binding affinity of SARS-CoV-2 S to ACE2 in Vero E6 cell. Consistent with the findings from the confocal imaging, full-length S, S1, and RBD, but not S2, showed high binding affinity to ACE2. Furthermore, the western blot analysis demonstrated that EGCG inhibited the binding of S (full-length S, S1 and RBD) to ACE2 in a dose-dependent manner (**Fig. 6E-6H**).

To validate the western blot results, we analyze the effect of EGCG on the binding of the S to ACE2 by ELISA. We found that the recombinant RBD could bind to ACE2 with EC^50^ of 4.08 ng/mL, while EGCG significantly decreased the binding affinity of RBD (4.7-fold) with EC^50^ of 19.19 ng/mL (**Fig. 6I**). In addition, EGCG also diminished the binding affinity of full-length S to ACE2 by 2.5-fold (EC_50_: 43.48 to 107.6 ng/mL, **Fig. 6I**). The inhibitory effect of EGCG on RBD or full-length S binding to ACE2 was dose-dependent (**Fig. 6J**).

## Discussion

Our rationales to study the anti-SARS-CoV-2 activity of GTB and its major ingredients were based on the followings: **1**. Green tea is the most consumed beverage in Asian countries where the morbidity and mortality of COVID-19 are significantly lower than Europe, North and Latin America; **2**. Green tea and its major catechin EGCG have a broad-spectrum antiviral effect on some pathogenic human viruses, including RNA viruses and those causing respiratory diseases^[14-20]^; **3**. Studies from clinical trials indicate that the consumption of catechins-containing GTB can reduce influenza infection rates^[26, 27, 46]^; **4**. Green tea catechins have anti-inflammatory and anti-oxidative properties which are beneficial for reducing severity of COVID-19. In the present study, we demonstrated that GTB could inhibit both wildtype and variant SARS-CoV-2 S-mediated infection (**Fig. 2B and 2C**). The inhibitory effect on the virus infection was highly potent as GTB at a 320-fold dilution remained significantly effective (**Fig. 2B and 2C**).

We subsequently demonstrated that the major green tea catechin EGCG was highly effective in inhibiting live SARS-CoV-2 infection of human lung epithelial cells (Calu-3) and HEK293-hACE2 cells which was evidenced by decrease of viral RNA, N protein and virus-induced plaques (**Fig. 4A-4C**). The catechins are polyphenolic compounds contained in green tea and contribute to most of its biological effects. EGCG is considered as the representative tea catechin with the highest activity^[40]^. Among 4 green tea catechins at noncytotoxic doses, EGCG was the most potent in inhibiting infectivity of the pseudovirus bearing WT S of SARS-CoV-2 (**Fig. 3A**). EGCG also significantly suppressed the infectivity of pseudoviruses bearing the S with mutations of the newly emerged variants (UK-B.1.17, SA-B.1.35, and CA-B.1.429) (**Fig. 3C and 3D**). In addition, among the catechins, EGCG was the strongest inhibitor of live human coronavirus (HCoV OC43) infection (**Fig. 4D and 4E**), which agrees with a recent report that EGCG treatment decreased the levels of HCoV OC43 viral RNA and protein in infected cell media through inhibition of 3CL-protease activity of the virus^[47]^. These data provide the direct evidence to support the assumption that high green tea consumption was associated with low morbidity and mortality of COVID-19 at the level of individual countries^[30]^.

To determine the mechanisms of anti-SARS-CoV-2 action by EGCG, we examined the impact of EGCG on the interaction between ACE2 and the RBD of SARS-CoV-2 S. We found that EGCG blocked viral attachment to host cells through interfering with the engagement of the SARS-CoV-2 S and ACE2 receptor (**Fig. 6A, 6E and 6F**). Our further studies showed that EGCG could block the binding of the RBD of S to ACE2 receptor (**Fig. 6D, 6E and 6H**). These findings suggest that the interaction of EGCG with the virion surface proteins prevents the viral attachment to the host cells. This suggestion is also supported by the observation that EGCG was the most effective when it was premixed with the virus prior to infection (**Fig. 5A**). A most recent study^[48]^ reported that teas and their constituents could inactivate SARS-CoV-2 *in vitro* through their interaction with the virions rather than cells because pretreatment of virion but not cells with catechin derivatives significantly suppressed the viral infection. The studies of EGCG effect on different viruses showed that EGCG inhibits viruses by a common mechanism: EGCG acts directly on the virion surface proteins without affecting the fluidity or integrity of the virion envelopes^[18]^. EGCG blocks the primary low-affinity attachment of virions to cells, because EGCG competes with heparan sulfate or sialic acid moieties in cellular glycans for virion binding^[18]^. Therefore, there is a potential for developing EGCG as a potent entry inhibitor of SARS CoV-2 infection.

During the preparation of this manuscript, two independent research groups reported in non-peer-reviewed preprints ^[48-50]^ that green tea has antiviral activity against SARS-CoV-2. Frank *et al*. showed that one-minute incubation of live SARS-CoV-2 with GTB resulted in ∼80% decrease of the viral titer^[49]^.

Ohgitani et al revealed that both black and green tea could inactivate SARS-CoV-2 *in vitro* due to their inhibitory effect on interaction between RBD of S protein and ACE2^[48]^. While these independent and preliminary reports support our finding that green tea has anti-SARS-CoV-2 activity, it remains to be determined whether green tea or its ingredients have the ability to inhibit the new variants of SARS-CoV-2. We now demonstrate for the first time that both GTB and EGCG could significantly inhibit, in a dose-dependent manner, the infectivity mediated by mutant S proteins of the newly emerged SARS-CoV-2 variants (UK-B.1.1.7, SA-B.1.351, and CA-B.1.429). This finding is clinically important, as SARS-CoV-2 frequently undergoes mutations, especially within its spike protein.

Clinical studies documented that EGCG was safe and well tolerated by the study subjects^[51, 52]^. Purified EGCG at the doses up to 1600 mg daily was well-tolerated and safe to use clinically^[53]^. Oral use of EGCG in mouthwash formulation at the dose of 35-87 mM for 2 min daily for 7 days had little side effects^[54]^. The FDA approved drug formulation of EGCG, Polyphenon E ointment (Veregen) for topical treatment of genital warts, has little adverse effects ^[55, 56]^. Our evaluation of cytotoxicity in the cell cultures showed that the treatment with GTB at dilution up to 20-fold or EGCG at dose up to 100 μM for 72 h had no cytotoxicity to all four cell types used in this study (**Supplemental Fig. 3**). These observations are consistent with the reports by others^[32, 57]^. As one of the most widely consumed beverages in Asian countries, green tea is used by many people at different ages on a daily basis. Therefore, oral rinsing and gargling with GTB should be safe and well suitable for pre- and postexposure prophylaxis against SARS-CoV-2. It would be of great epidemiological interest to determine whether green tea consumption does have negative impact on morbidity and mortality of COVID-19.

Because nasal and oropharyngeal mucosa are the primary sites for the initial infection of SARS-CoV-2, protecting these areas is crucial for preventing the viral transmission/dissemination, and development of COVID-19. Therefore, it is important to evaluate whether nasal and oral mucosal preexposure (oral rinse or gargling) of GTB or EGCG can prevent SARS-CoV-2 transmission and infection. It has been reported that gargling tea could lower the incidence of influenza virus infection^[58, 59]^. A prospective cohort study on the effect of green tea gargling in elderly residents (>65 yr.) living in a nursing home showed that the incidence of influenza infection decreased compared to gargling with water^[59]^. By extrapolating, it is likely that the same benefits of EGCG shown for decreasing influenza infection are applicable for the prevention of SARS-CoV-2 transmission/infection. This supposition, however, needs to be demonstrated in appropriate animal models and clinical studies. In addition, it is important to develop EGCG formulations that have stable, effective, and long-lasting action on inactivation of SARS-COV-2 and its variants. It should be noted that a clinical trial to evaluate the effects of an oral formulation of EGCG (Previfenon®) in the prevention of COVID-19 in health care workers was recently proposed (https://clinicaltrials.gov/ct2/show/NCT04446065).

Taken together, our data as well as the findings just reported by others^[48-50]^ indicate the possibility that consumption of green tea or its active ingredient EGCG is beneficial for preventing or reducing SARS-CoV-2 transmission and infection. However, clinical significance and application of ours and the reported *in vitro* findings need to be validated in future investigations, particularly in clinical studies of the anti-SARS-CoV-2 effects of EGCG as a viral entry inhibitor. Nonetheless, given its low toxicity, anti-inflammation, antioxidant, and anti-SARS-CoV-2 variant properties, use of GTB or EGCG is likely to minimize the SARS-CoV-2 spread, ameliorate symptoms and disease severity.

## Author contributions

J.B.L., F.Z.M., P.W. Z.Y.W. and L.N.Z. performed and analyzed pseudovirus assays. B.H.B, S.S and W.H. designed and construct spike variant plasmids. A.K. and J.J.L. obtained and characterized clinical samples. G.X.L, T.W., S.C.L. and Q.L. performed live SARS-CoV-2 assays. J.G.W., Q.L., W.H. and W.Z.H. supervised the research and revised manuscript. J.B.L and X.W. wrote the initial draft, with the other authors providing editorial comments.

## Acknowledgements

This study was supported by Temple University seed grant (142398 and 160979). We thank Peihui Wang (Shandong University) for providing pCAG-SARS-CoV-2-S and pCAG-hACE2-Flag plasmids, Fang Li (University of Minnesota) for providing pcDNA3.1-SARS2-S (codon-optimized). The following reagent was deposited by the Centers for Disease Control and Prevention and obtained through BEI Resources, NIAID, NIH: SARS-Related Coronavirus 2, Isolate USA-WA1/2020, NR-52281; Human Coronavirus, OC43, NR-52725; Quantitative PCR (qPCR) Control RNA from Heat-Inactivated SARS-Related Coronavirus 2, Isolate USA-WA1/2020, NR 52347.

---

**Supplemental Figure 1.**
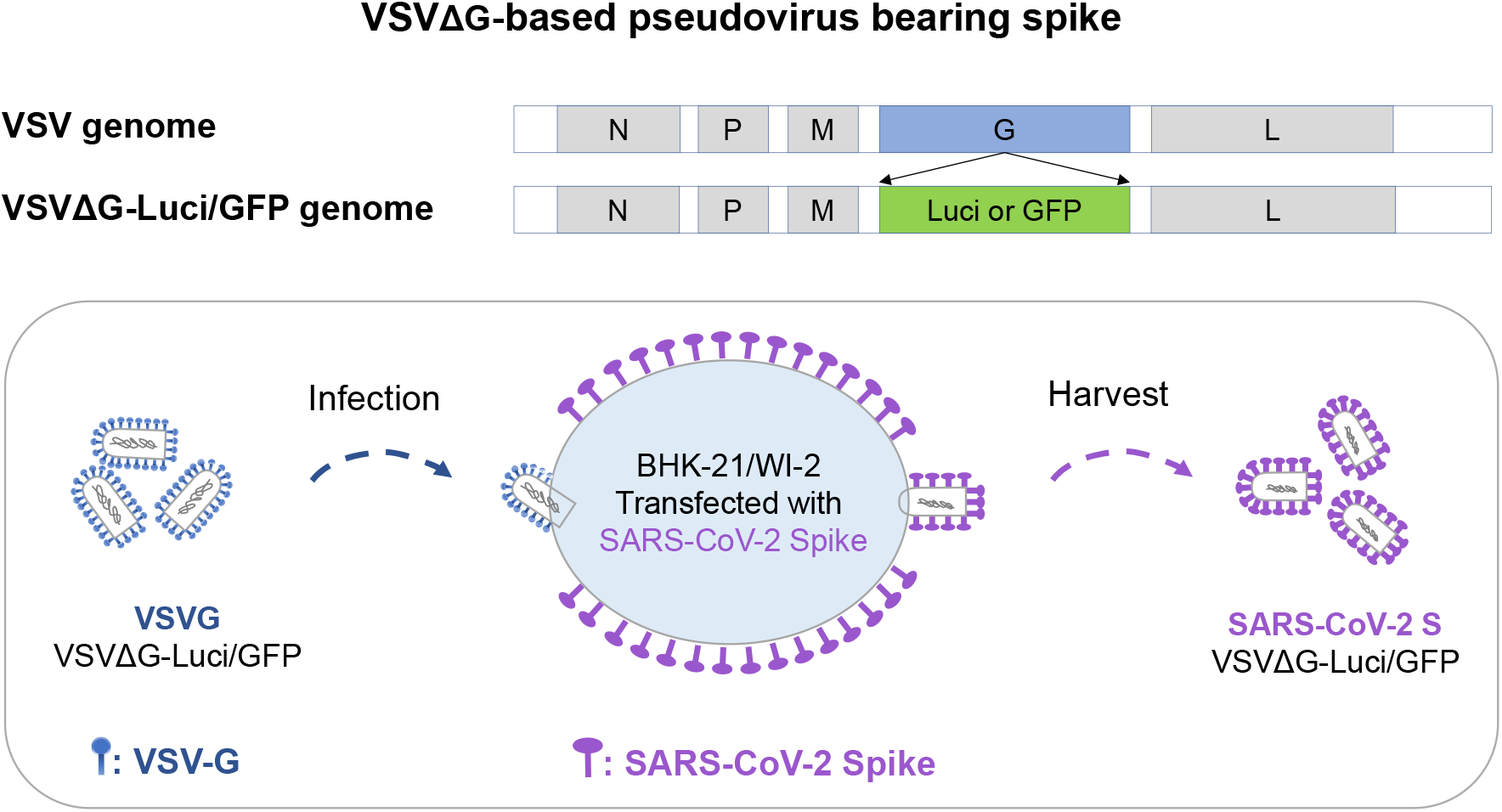
Generation of VSVΔG-based pseudovirus bearing SARS-CoV-2 S protein (SARS-CoV-2 pseudovirus). BHK-21/WI-2 cells were transfected with pCAG-SARS-CoV-2 S plasmids 24 h prior to VSVΔG-Luci/GFP-based pseudovirus bearing VSVG (VSVG-VSVΔG-Luci/GFP) infection (MOI=4). The cells were washed with PBS for 3 times to remove unbounded VSVG-VSVΔG-Luci/GFP at 2 h postinfection, and cultured in 5% DMEM for 24 h. Virions-containing supernatant was harvested and centrifuged at 500 g for 5 min, filtered with 0.45 μm polyethersulfone membrane.

**Supplemental Figures 2.**
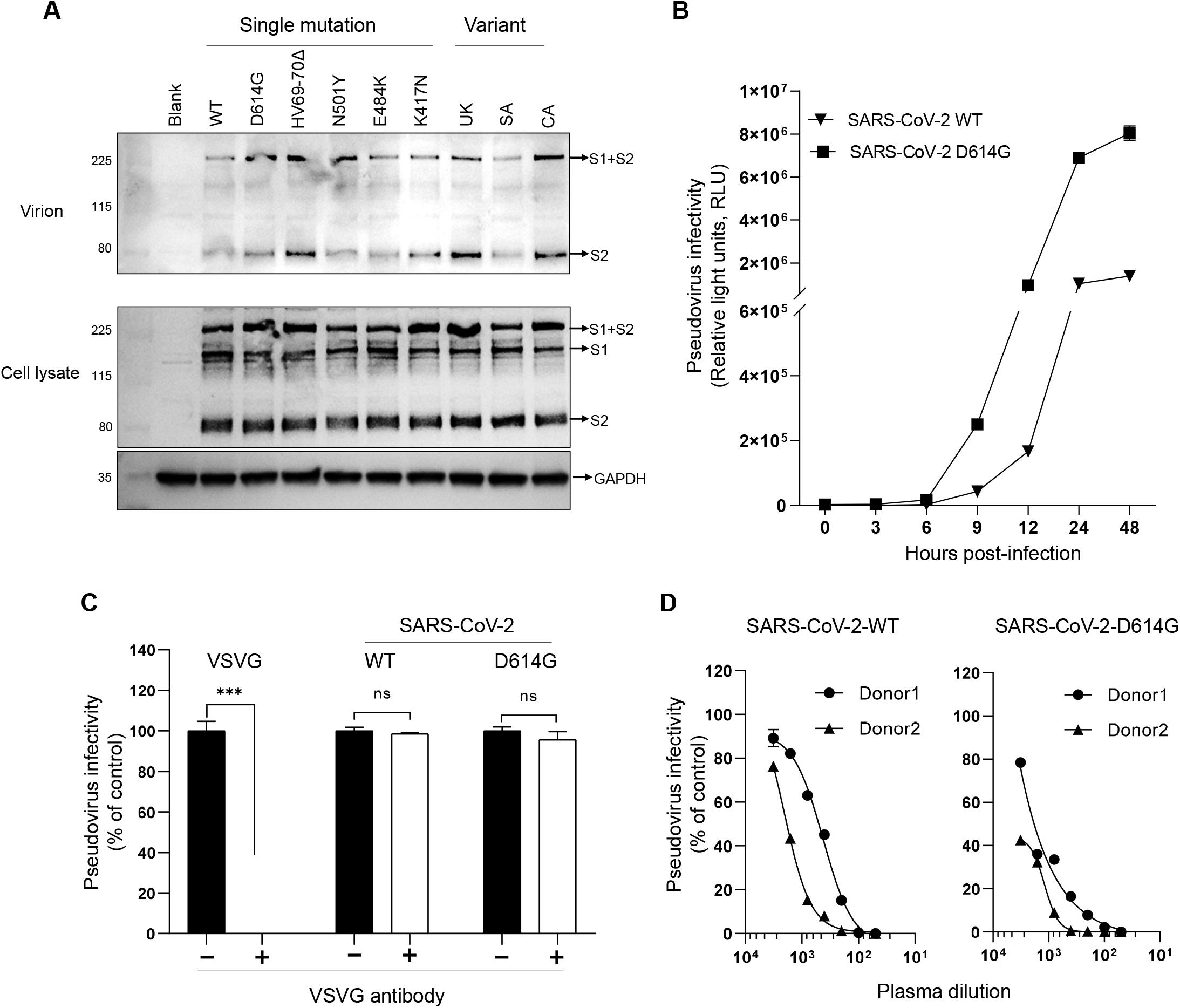
Verification of the incorporated SARS-CoV-2 spike in the pseudovirus. (**A**) Virion-containing supernatant and producer cells (BHK-21/WI-2) were harvested and subjected to western blot using SARS-CoV-2 convalescent plasma as the detection antibody (1:500 dilution). GAPDH was used as a loading control of western blot. (**B**) HEK239T-hACE2 cells were infected with VSVΔG-based pseudovirus bearing SARS-CoV-2 spike (SARS-CoV-2 pseudovirus) for indicated time, cells were lysed and subjected to luciferase activity detection. (**C**) The specificity of SARS-CoV-2 pseudovirus infection was tested against VSVG-specific antibody using VSVG-VSVΔG-Luci pseudovirus as the control. The data represent percentage infectivity relative to that of the untreated group. (**D**) Plasma samples from 2 convalescent patients of COVID-19 were serially diluted and incubated for 1 h at 37°C with indicated SARS-CoV-2 pseudoviruses and infectivity in HEK293T-hACE2 cells was then measured. The data represent percentage infectivity relative to that of the control (untreated). \*\*\**P*<0.001.

**Supplemental Figures 3.**
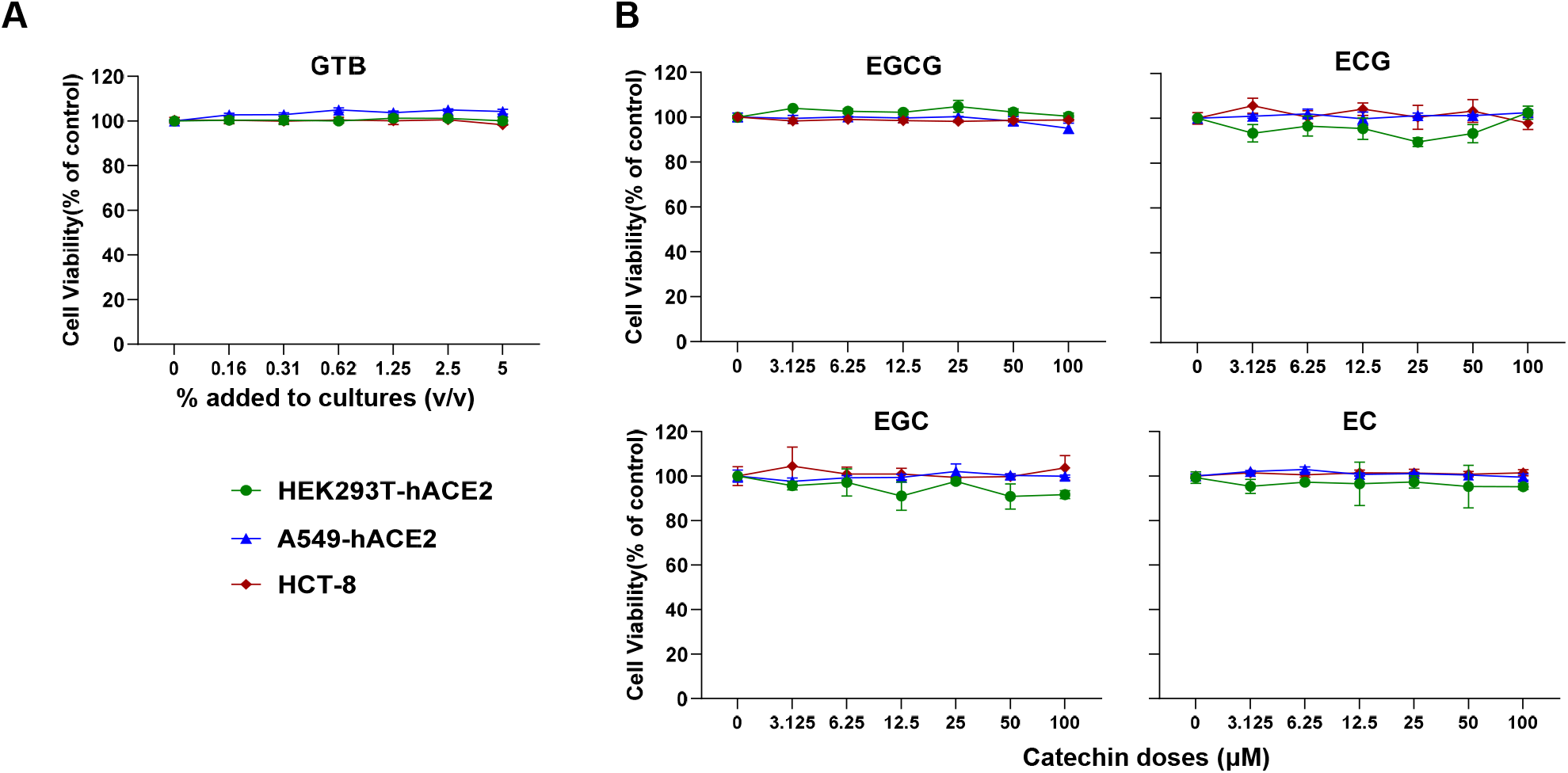
Green tea beverage (GTB) and the catechins have little cytotoxicity on the cells. (**A, B**) Cells (HEK293T-hACE2, A549-hACE2 and HCT-8) were treated with GTB or the catechins (EGCG, ECG, EGC and EC) at the indicated doses for 72h. The cell viability was assessed by MTS assay. Data represent percentage changes of the absorbance (490 nm) relative to untreated control.

**Supplemental Figures 4.**
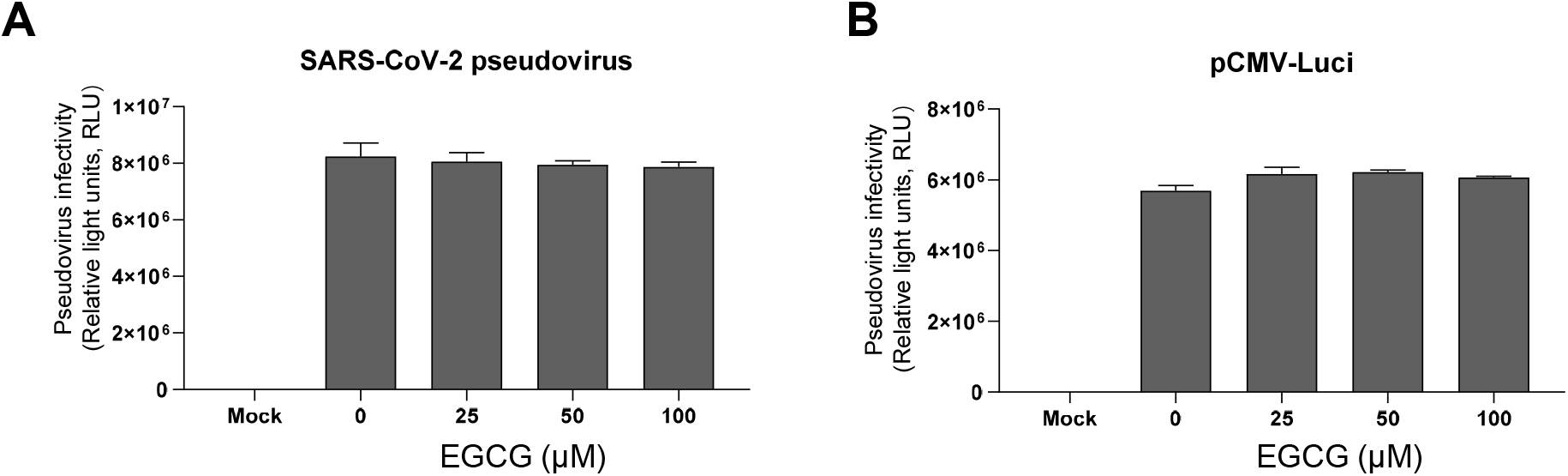
EGCG has no effect on luciferase activity and luciferase gene expression. **A**. HEK293T-hACE2 cells were infected with VSVΔG-based pseudovirus bearing SARS-CoV-2 S (SARS-CoV-2 pseudovirus) for 48 h, and then lysed, different dose of EGCG was added in the cell lysate when measuring luciferase activity. **B**. HEK293T-hACE2 cells were transfected with plasmid encoding luciferase, then treated with or without EGCG (0∼100 μM) for 48 h. Cells were lysed and subjected to luciferase activity detection.

**Table S1.**
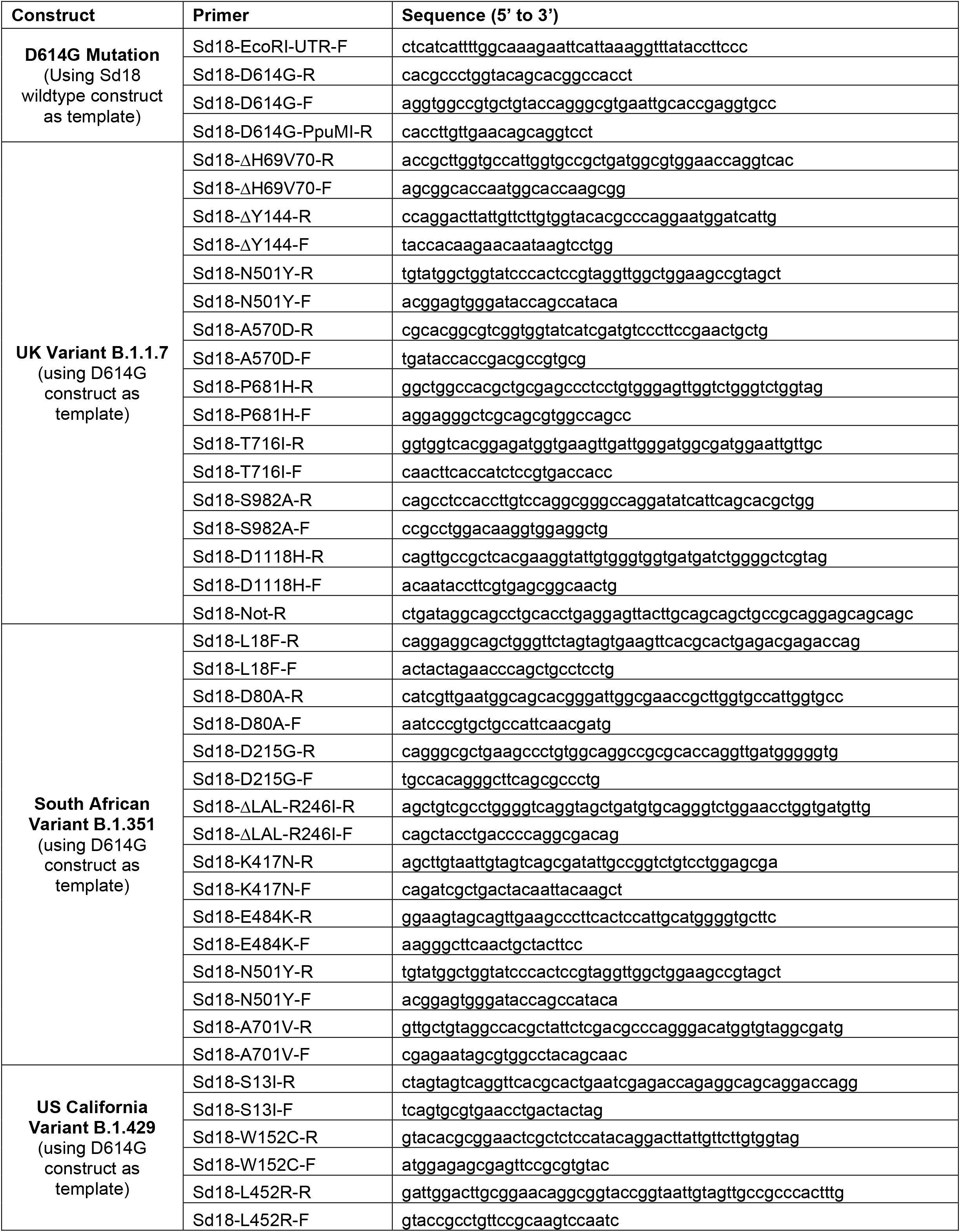
Mutagenic primers for each mutant of full-set S variants.

